# Spatial modulation of facial expression: enhanced recognition of faces behind the observer

**DOI:** 10.1101/2024.05.08.593080

**Authors:** Hideki Tamura, Yugo Kobayashi, Shigeki Nakauchi, Tetsuto Minami

## Abstract

Our ability to recognize facial expressions is crucial for understanding others’ emotions^1^ and facilitating smooth communication^2^. Numerous studies have explored how we perceive these cues, considering factors such as health^3^, social signals^4^ and personality traits^5^. However, most of this research involves observers facing a monitor and assessing facial stimuli presented directly in front of them. Real-life scenarios offer more diverse spatial dynamics, such as conversing with someone at a table or glancing back at a passerby. Thus, faces behind the observer might trigger heightened recognition, akin to reacting swiftly to a perceived threat. Herein, we demonstrate that facial expression recognition is influenced by spatial relationships, i.e., faces in front of versus behind the observer. Participants judged the expressions of faces appearing in front of or behind them in virtual space. The findings of three experiments reveal an enhanced level of recognition for faces behind the participant. Interestingly, this effect varies with emotional valence; anger is amplified merely by the presence of a face behind the observer, while happiness requires actively turning to the rear for enhancement to occur. These findings suggest a biological instinct for perceiving threats behind us, potentially influencing subsequent actions. Hence, spatial relationships may modulate facial expression recognition.

## Main

Participants, wearing head-mounted displays, viewed 3D computer-generated faces in virtual space (Figure 1A). These faces were transformed from neutral to angry or happy expressions using FaceGen software (see Supplement Information). In Experiment 1, participants directly observed faces either in front of them or 180 degrees behind them (Figure 1B upper row) and indicated whether each face appeared neutral or expressed anger/happiness. Psychometric functions were used to analyse their response probabilities, yielding the point of subjective equality (PSE) and just noticeable difference (JND) for comparison across conditions (Figure 1C). Regardless of the target expression, faces were perceived as more intense in the behind condition (Figure 1D and 1E). This finding suggests enhanced facial expression recognition when turning to the rear. While task-related factors might influence this outcome, with a larger JND for anger in the behind condition (Figure 1F), happiness discrimination showed no difference between the front and rear conditions (Figure 1G). Thus, task characteristics alone cannot explain these results.

**Figure 1.**
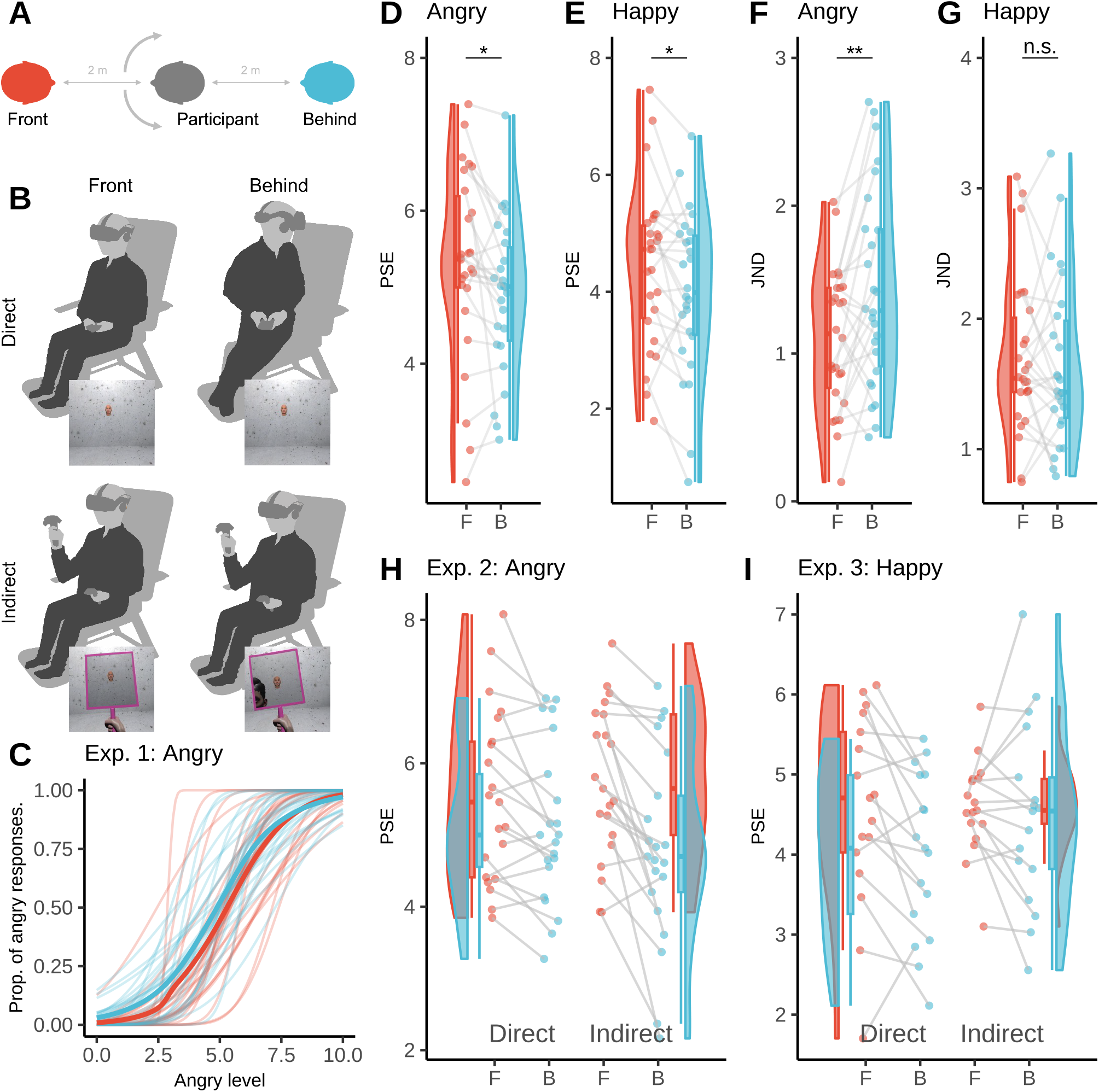
Experimental setup, observation conditions, and experimental results. (A) Spatial relationship between the participant and face stimulus. (B) Experimental conditions with the observation factor (direct/indirect in row) and direction factor (front/behind in column). Experiment 1 included only the directly in front and directly behind conditions. Experiments 2 and 3 included all four conditions. The bottom image in each cell is an example of the participant’s view. (C) Psychometric function from the participants’ responses in the angry condition in Experiment 1 as an example. The horizontal axis indicates the angry levels of the stimuli (0: neutral, 10: angry). The vertical axis indicates the proportions of angry responses. The thick and thin curves illustrate the average of the participants and individual data, respectively. (D-G) The PSEs and JNDs of the angry and happy conditions in Experiment 1. The horizontal axis indicates each direction (F: front, B: behind). The vertical axis indicates the PSE (D and E) and JND (F and G). The raincloud plot shows the individual data, box plot, and their distributions. For PSEs, lower values indicate that the participants were more likely to respond to the facial stimulus as the target expression (angry/happy). Asterisks indicate significant differences: ^*^*p <* .05, ^* *^ *p <* .01. The behind condition was perceived as angrier (D) and happier (E) than the front condition (Angry: *M*_*D*_ = 0.42, 95% CI [0.09, 0.75], *t*(25) = 2.66, *p* = .014; Happy: *M*_*D*_ = 0.43, 95% CI [0.05, 0.81], *t*(25) = 2.35, *p* = .027). In contrast, while it was more difficult to judge the angry facial expression in the behind condition (*M*_*D*_ = *−*0.33, 95% CI [*−*0.54, *−*0.12], *t*(25) = *−*3.28, *p* = .003), there was no difference in the happy facial expression (*V* = 198.00, *p* = .582). (H) The PSEs of Experiment 2 (angry). There was a main effect in the direction condition (*F* (1, 20) = 36.36,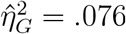 .076, 90% CI [.000, .302]), as well as a significant interaction (*F* (1, 20) = 4.75,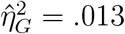 .013, 90% CI [.000, .184]). The direct observation of the behind face demonstrated an enhanced perception of angry than that for the front condition (Δ*M* = 0.39, 95% CI_Tukey(4)_ [0.05, 0.74], *t*(20) = 3.18, *p*_Tukey(4)_ = .023), supporting the results of Experiment 1 and revealing which effect was stronger in the indirect observation (Δ*M* = 0.93, 95% CI_Tukey(4)_ [0.38, 1.48], *t*(20) = 4.71, *p*_Tukey(4)_ *<* .001). (I) The PSEs of Experiment 3 (happy). There was a significant interaction (*F* (1, 16) = 4.76, *p* = .044, 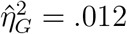, 90% CI [.000, .204]). We also found the same enhancement of direct observation for the behind face in the happy condition as well (Δ*M* = 0.56, 95% CI_Tukey(4)_ [0.04, 1.08], *t*(16) = 3.06, *p*_Tukey(4)_ = .034), but this did not hold for indirect observation (Δ*M* = 0.11, 95% CI_Tukey(4)_ [*−*0.51, 0.73], *t*(16) = 0.52, *p*_Tukey(4)_ = .952). In (H-I), the formats were the same as those used in (D-G).

In the previously mentioned behind condition, it remained unclear whether the enhancement in expression perception was due to physical turning around or simply the presence of the face behind the observer. To address this issue, we explored the effects of conditions in which participants could judge faces behind them without physically turning. Participants utilized a “virtual hand-mirror” via controller input to reflect faces behind them (mirror; Figure 1B bottom right), and a “virtual glass tool” that resembled the hand-mirror but showed faces directly in front, akin to viewing through a glass pane (glass; Figure 1B bottom left). In Experiment 2 (Figure 1H), which focused on anger as the target expression, the results showed a significant effect of front/behind on PSE, with faces being perceived as angrier when behind, regardless of physical turning. Moreover, there was an interaction that indicated a greater difference in expression intensity perception between the front and behind faces in the tool-operated condition without physical turning. Experiment 3 (Figure 1I) investigated happiness as the target expression, replicating Experiment 1’s findings with bodily movement resulting in rear faces being perceived as happier. However, using the tool showed no difference in expression perception between the front and behind faces. These results suggest that the perception of rear face expression varies based on the emotional valence of the observed face. Notably, there were no significant JNDs found across conditions in Experiments 2 or 3, indicating that expression perception differences are not solely explained by observation conditions (Figure S1). These findings imply the involvement of a specialized expression recognition mechanism for faces behind the observer.

The current findings demonstrate that spatial orientation relative to the observed face influences facial expression recognition. Through the integration of visual and somatic sensory inputs, we discern the spatial dynamics between ourselves and others, prompting a specialized recognition process for faces positioned behind us, where visual information retrieval is more challenging. Stimuli that are positioned behind us inherently pose visual perceptual difficulties compared to those in front, leading to partial or peripheral information acquisition^6,7^. Thus, heightened expression recognition in these scenarios may serve as a compensatory mechanism. Moreover, we observed an additional compensatory effect tied to the emotional valence of the stimulus. Particularly, for potentially threatening angry expressions, facial recognition was further intensified, with faces behind being perceived as angrier even without physical reorientation. This heightened perception of anger from behind may serve to avert potential threats, as indicated by findings suggesting greater comfortable interpersonal distances^8^, especially for rear spaces^9^. This augmented perception of threat-related expression could stem from an innate survival strategy, conferring an evolutionary advantage^10^. In essence, the visual system appears to recalibrate expression recognition, considering both the spatial relationship between observer and face and the perceived threat level conveyed by the facial expression itself.

The phenomenon revealed in this study likely involves a complex interplay of factors encompassing expression processing, bodily sensation, and the estimation of interpersonal distances. To comprehensively understand visual processing mechanisms in real-life contexts, it is imperative to explore visual stimuli presentation under conditions that offer greater degrees of freedom that extend beyond frontal views. Moreover, beyond facial expression recognition alone, the perception of diverse stimulus modalities may also vary based on spatial relationships with stimuli. Alternatively, this observed phenomenon could underscore the significance of faces and interpersonal spatial dynamics, warranting further investigation.

In summary, our findings demonstrate enhanced facial expression recognition for stimuli positioned behind the observer. Importantly, this enhancement occurs even without the need for physical reorientation, suggesting more efficient visual information processing by the visual system.

## Acknowledgements

This work was supported by JSPS KAKENHI (Grant Numbers JP22K17987, JP20H05956, and JP20H04273). This study was based on the results obtained from project JPNP20004, which was subsidized by the New Energy and Industrial Technology Development Organization (NEDO). The authors wish to thank Teruyuki Inoue for supporting the data collection.

## Data Availability

The code and data underlying the results presented in the study are available from the Open Science Framework repository (https://osf.io/zpuey/).

## Declaration of generative AI and AI-assisted technologies in the writing process

During the preparation of this work the authors used ChatGPT 3.5 in order to improve language, and the manuscript has been proofread by native English speakers through an English proofreading service. After using the tool and service, the authors reviewed and edited the content as needed and take full responsibility for the content of the publication.

## Supplemental Information: Supplemental figures

Spatial modulation of facial expression: enhanced recognition of faces behind the observer

Hideki Tamura, Yugo Kobayashi, Shigeki Nakauchi, Tetsuto Minami

## Supplemental Information: Experimental Procedure

### Participants

We recruited 34, 24, and 24 participants, who were students from Toyohashi University of Technology, for Experiments 1, 2, and 3, respectively. The sample size of Experiment 1 (n = 34) was determined by performing a power analysis using G*Power (3.1.9.6)^1^ for the t test, with an effect size = 0.5, alpha = .05, and power = 0.8. The sample sizes of Experiments 2 and 3 (n = 24) were also determined by the same procedure for a within-participants two-factor analysis of variance interaction with the observation condition (direct or indirect) and the direction condition (front or behind), whose parameters were as follows: effect size = 0.25, alpha = .05, and power = 0.8. The average and standard deviation of the participants’ ages were 22.2 ± 1.5 (3 females and 31 males in Experiment 1), 23.1 ± 1.3 (3 females and 21 males in Experiment 2), and 22.7 ± 1.1 (4 females and 20 males in Experiment 3). In the process of recruiting participants for the experiment, individuals of any gender were recruited. The Committee for Human Research of the Toyohashi University of Technology approved the experimental procedures, which were conducted in accordance with the Declaration of Helsinki. Written informed consent was obtained from all participants after the procedural details were explained to them.

### Apparatus

The stimuli were presented on a head-mounted display (HMD: HTC VIVE Cosmos, 1440 x 1700 resolution per eye, 90 Hz refresh rate, 110-degree field of view) with a Windows 10 desktop computer equipped with a Core i7-11700 and 32 GB RAM and a graphic board (NVIDIA GeForce RTX 3070). We ran the experiment using Unity 2021.1.25f1 and Steam VR 2.3.3 with an additional asset, Playmaker 1.9.7. To obtain the participants’ responses, a numerical keypad (Experiment 1) or VIVE Cosmos Controller (Experiments 2 and 3) was used.

### Stimuli

We generated a 3D human face model using FaceGen Modeller Core 3.29 (Singular Inversions Inc.). We randomly created two female and two male individual faces with the “East Asian racial group” setting. For the angry condition, each model’s facial expression was changed by the “Expression Anger” parameter from 0 (neutral) to 3, 4, 5, 6, 7, or 10 (angry) to represent the seven expression levels. In addition, we modified the “Phoneme ee” parameter, which is related to the degree of mouth openness of the model, from 10 (neutral) to 7, 6, 5, 4, 3, or 0 (angry). When this parameter decreases, the model gradually closes its mouth. This was done to prevent the judgement of expressions solely based on the degree of mouth opening. The facial models for the happy condition were also created in the same way except that the “Expression SmileOpen” parameter was changed. In addition to changing the parameters of facial expression, we also changed “Phoneme ee,” which reflects the shape of the model’s mouth. The number of stimuli used was 56 (4 individual faces x 7 facial expression levels x 2 emotions). Experiment 1 used stimuli of both emotions (angry and happy), and Experiments 2 and 3 used stimuli of only one emotion, i.e., angry and happy, respectively. Note that participants recognized the facial expressions modified by FaceGen as intended expressions^2,3^.

### Procedure

**Experiment 1**. Participants wearing an HMD took a seat on a chair placed in the centre of the experiment room and held a keypad with both hands. The position where the participant sat comfortably and the height of the participant’s head were set as the origin. In the virtual environment, the participants were surrounded by walls with textures^4^. Each trial was conducted according to the following procedure (see Figure S2). First, participants were instructed to return their head position to the origin. Next, a face model appeared either in front or behind. A fixation point (in the front condition) or an arrow pointing left/right (in the behind condition) was presented at the front. In the front condition, after participants fixated on the fixation point for 500 ms, a stimulus (0.28 × 0.19 m; 8.00 × 5.44 degrees) was presented at a position located 2 metres away from the participant in front.

After the stimulus remained in the participant’s field of view for 2,000 ms, the response phase started, during which the participants were asked to indicate whether the stimulus appeared neutral or angry using the keypad. When the target facial expression was happy, the choices were either neutral or happy. In the behind condition, a stimulus was presented 2 metres away at a position located 180 degrees behind, and participants were required to rotate their bodies according to the direction of the presented arrow to see the stimulus for 2,000 ms. After the response was accepted, the next trial began. For each type of stimulus, the front condition consisted of 2 repetitions with 2 trials each, while the behind condition consisted of 1 repetition with 2 trials each in both the left and right directions. Thus, each participant completed a total of 224 trials divided into 4 blocks, with each expression condition occurring twice (4 identities x 7 levels x 2 trials x 2 front/back x 2 expressions). The order of running for the angry and happy expression blocks was counterbalanced across participants. Participants were allowed to take breaks of any duration between blocks. The correspondence between the two response expressions and the two response keys was also counterbalanced across participants.

**Experiment 2**. The basic setup was similar to that used in Experiment 1, with the difference that participants held controllers in both hands instead of a keypad and responded using either trigger. The direct front and direct behind conditions were the same as those described for Experiment 1. In the indirect front condition, a fixation point was presented, and after participants fixated on it for 500 ms, text indicating which hand held the virtual tool (left or right) was displayed for 1,000 ms. Subsequently, the fixation point reappeared, and after being contained within the frame of a virtual glass for 1,000 ms, a stimulus was presented at that position. After the stimulus was observed for 2,000 ms, the response phase started. In the indirect behind condition, a face stimulus was presented 180 degrees behind, and simultaneously, text indicating which hand held the virtual mirror (left or right) was displayed for 1,000 ms. Participants saw the stimulus through the mirror and entered the response phase after observing it for 2,000 seconds. The response phase was the same as that described for Experiment 1, with the target expression being angry. Each participant completed 224 trials in random order (4 identities x 7 expressions x 2 left/right hands x 2 front/behind x 2 observations). The experiment was divided into 4 blocks, with breaks of any duration allowed between blocks. The correspondence between the two response expressions and the two response triggers was counterbalanced across participants.

Additionally, the stimuli in the indirect conditions were presented to match the size of those in the direct conditions to prevent any influence of stimulus size on expression recognition. Furthermore, to increase the sense of body ownership, either a male or female avatar representing the participant, depending on the participant’s reported gender, was displayed on the participant’s position. Note that we adjusted the avatar’s position towards the lateral direction to easily fit its body within the frame of the virtual mirror.

**Experiment 3**. The target expression was changed to “happy,” and the stimuli and responses were adjusted accordingly for the happy condition. Otherwise, the setup remained the same as that used in Experiment 2.

### Data analysis

We analysed the data and performed statistical tests using R 4.3.3. One of the participants in Experiment 1 was excluded from the analysis because the experiment terminated halfway due to an unexpected issue. The proportion of trials where participants responded with the target facial expression (angry/happy) was computed for each participant, condition, and expression level. We fitted the psychometric functions indicating the angry/happy proportion using a generalized linear model with a probit link function.

Then, PSEs and JNDs were determined for each participant and condition. We identified outliers from the data of the participants using the interquartile range (IQR) method (“identify_outliers” function) for the PSEs and JNDs. The outliers identified by any condition in each experiment were excluded from the subsequent analysis; there were 7, 3, and 7 participants excluded from Experiments 1, 2, and 3, respectively. In Experiment 1, we performed a t test or Wilcoxon test between the front and behind conditions for the PSEs and JNDs in each target facial expression (see Tables 1–4). In Experiments 2 and 3, we performed a two-way repeated-measures analysis of variance for the direction factor (front/behind) and observation factor (direct/indirect) for the PSEs (see Tables 5–7 in Experiment 2; Tables 8 and 9 in Experiment 3) and JNDs (see Figure S1). We performed the Shapiro–Wilk test to check the normality of the data in each condition and used the Tukey method for multiple comparisons. Note that even though the data of the indirect front conditions for the JNDs in Experiment 3 were not normally distributed according to the Shapiro–Wilk test (statistic = 0.87, p = 0.02), the quantile–quantile (QQ) plot showed that all the points fell approximately along the reference line. Thus, from this visual inspection, we assumed the normality of those data.

**Table 1.**
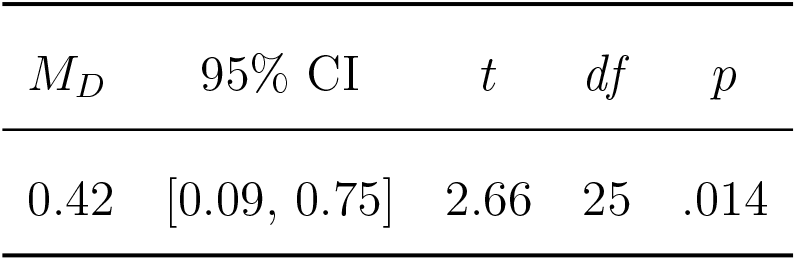
t test for PSEs (Angry)

**Table 2.** t test for PSEs (Happy)

| $M_D$ | 95% CI | $t$ | $df$ | $p$ |
| --- | --- | --- | --- | --- |
| 0.43 | [0.05, 0.81] | 2.35 | 25 | .027 |

**Table 3.**
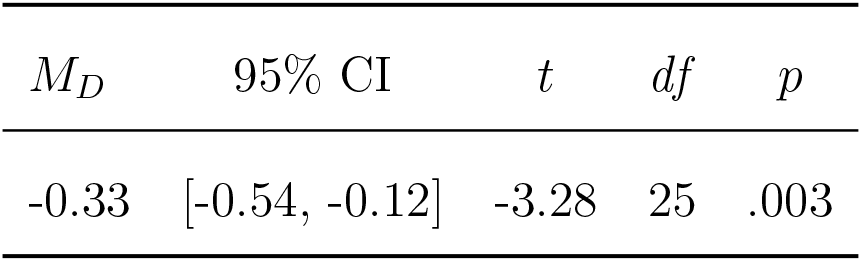
t test for JNDs (Angry)

| $M_D$ | 95% CI | $t$ | $df$ | $p$ |
| --- | --- | --- | --- | --- |
| -0.33 | [-0.54, -0.12] | -3.28 | 25 | .003 |

**Table 4.** Wilcoxon test for JNDs (Happy)

| $V$ | $p$ |
| --- | --- |
| 198.00 | .582 |

**Table 5.**
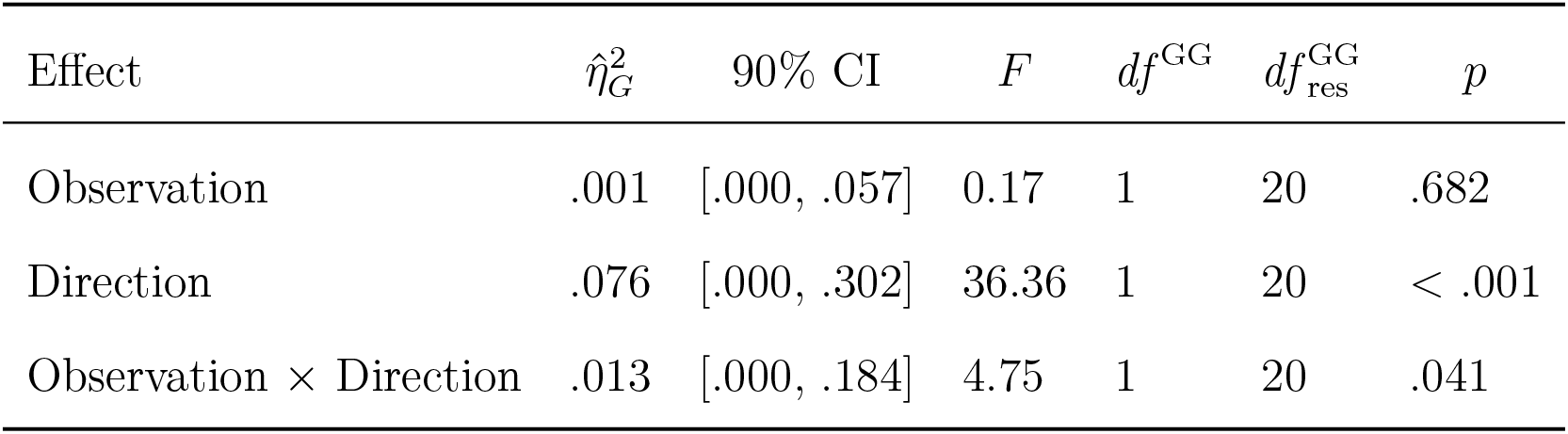
Summary of two-way repeated-measures ANOVA of PSEs in Experiment 2.

| Effect | $\hat{\eta}_G^2$ | 90% CI | $F$ | $df^{\text{GG}}$ | $df_{\text{res}}^{\text{GG}}$ | $p$ |
| --- | --- | --- | --- | --- | --- | --- |
| Observation | .001 | [.000, .057] | 0.17 | 1 | 20 | .682 |
| Direction | .076 | [.000, .302] | 36.36 | 1 | 20 | < .001 |
| Observation × Direction | .013 | [.000, .184] | 4.75 | 1 | 20 | .041 |

**Table 6.** Post hoc comparisons - Direction.

| Contrast | $\Delta M$ | 95% CI | $t$ | $df$ | $p$ |
| --- | --- | --- | --- | --- | --- |
| Front - Behind | 0.66 | [0.43, 0.89] | 6.03 | 20 | < .001 |

**Table 7.**
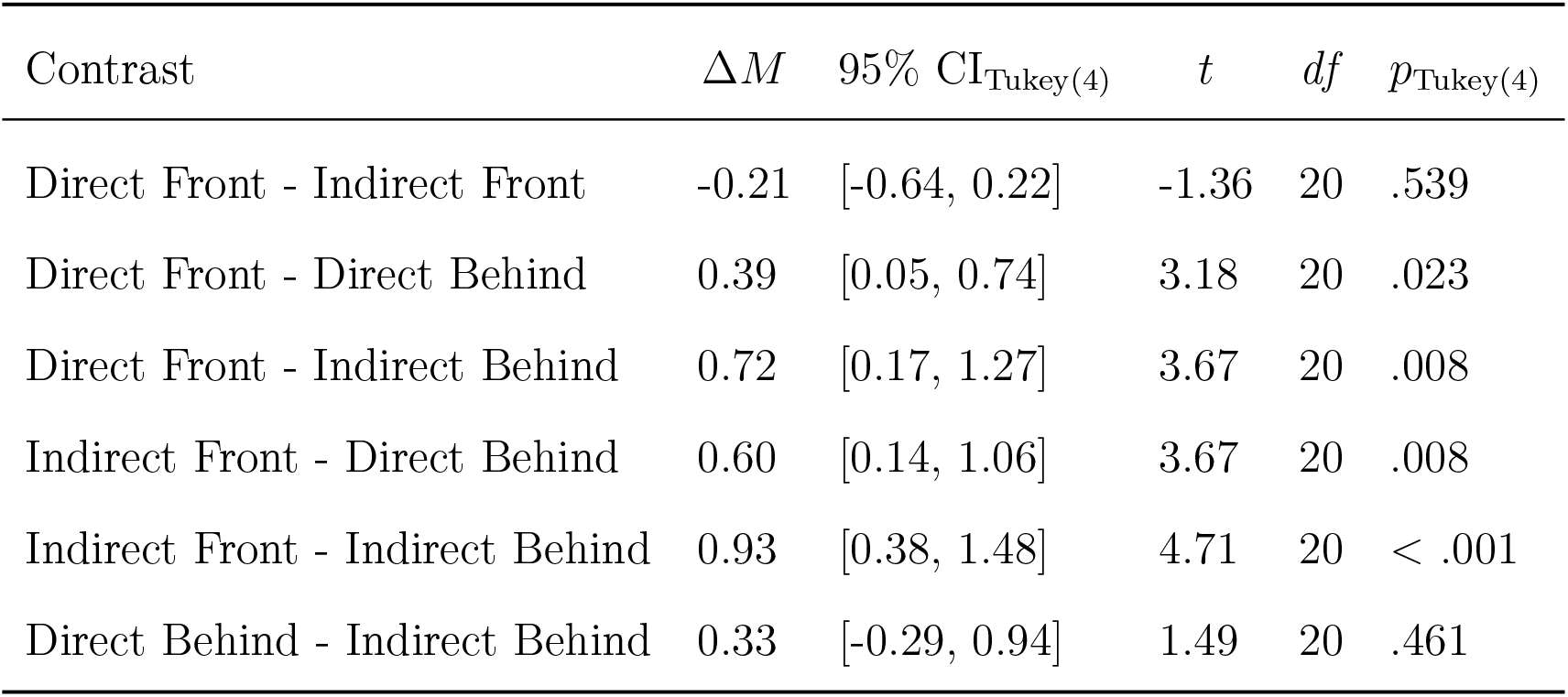
Post hoc comparisons - Observation * Direction.

| Contrast | $\Delta M$ | 95% CI<sub>Tukey(4)</sub> | $t$ | $df$ | $p_{\text{Tukey}(4)}$ |
| --- | --- | --- | --- | --- | --- |
| Direct Front - Indirect Front | -0.21 | [-0.64, 0.22] | -1.36 | 20 | .539 |
| Direct Front - Direct Behind | 0.39 | [0.05, 0.74] | 3.18 | 20 | .023 |
| Direct Front - Indirect Behind | 0.72 | [0.17, 1.27] | 3.67 | 20 | .008 |
| Indirect Front - Direct Behind | 0.60 | [0.14, 1.06] | 3.67 | 20 | .008 |
| Indirect Front - Indirect Behind | 0.93 | [0.38, 1.48] | 4.71 | 20 | < .001 |
| Direct Behind - Indirect Behind | 0.33 | [-0.29, 0.94] | 1.49 | 20 | .461 |

**Table 8.** Summary of two-way repeated-measures ANOVA of PSEs in Experiment 3.

| Effect | $\hat{\eta}_G^2$ | 90% CI | $F$ | $df^{\text{GG}}$ | $df_{\text{res}}^{\text{GG}}$ | $p$ |
| --- | --- | --- | --- | --- | --- | --- |
| Observation | .015 | [.000, .213] | 2.14 | 1 | 16 | .163 |
| Direction | .028 | [.000, .250] | 3.78 | 1 | 16 | .070 |
| Observation × Direction | .012 | [.000, .204] | 4.76 | 1 | 16 | .044 |

**Table 9.** Post hoc comparisons - Observation * Direction.

| Contrast | $\Delta M$ | 95% CI<sub>Tukey(4)</sub> | $t$ | $df$ | $p_{\text{Tukey}(4)}$ |
| --- | --- | --- | --- | --- | --- |
| Direct Front - Indirect Front | -0.02 | [-0.59, 0.55] | -0.10 | 16 | > .999 |
| Direct Front - Direct Behind | 0.56 | [0.04, 1.08] | 3.06 | 16 | .034 |
| Direct Front - Indirect Behind | 0.09 | [-0.70, 0.88] | 0.34 | 16 | .986 |
| Indirect Front - Direct Behind | 0.58 | [0.02, 1.14] | 2.96 | 16 | .041 |
| Indirect Front - Indirect Behind | 0.11 | [-0.51, 0.73] | 0.52 | 16 | .952 |
| Direct Behind - Indirect Behind | -0.46 | [-1.01, 0.08] | -2.44 | 16 | .110 |

---

**Figure S1.**
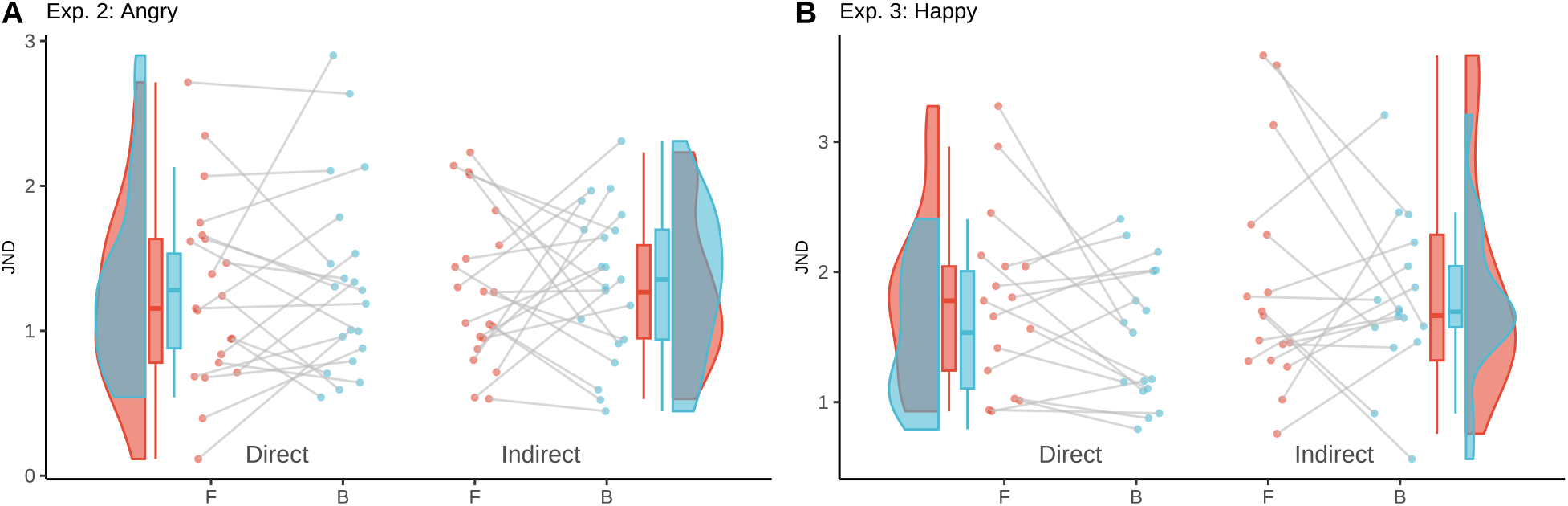
The JNDs of Experiments 2 and 3. For the JNDs, there were no main effects on the observation (direct/indirect) or direction (front/behind) conditions or interaction, regardless of the target facial expression, respectively: (A) Angry in Experiment 2 (*F* (1, 20) = 0.08, 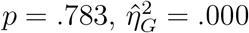, 90% CI [.000, .043], *F* (1, 20) = 0.46, *p* = .507, 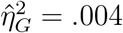, 90% CI [.000, .127], *F* (1, 20) = 0.04, 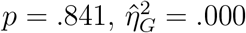, 90% CI [.000, .020]). (B) Happy in Experiment 3 (*F* (1, 16) = 1.55, 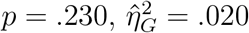, 90% CI [.000, .231], *F* (1, 16) = 1.77, *p* = .202, 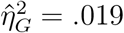, 90% CI [.000, .228], *F* (1, 16) = 0.30, *p* = .594, 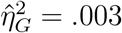, 90% CI [.000, .140]). The formats were the same as those in described for Figure 1 (D-G).

**Figure S2.**
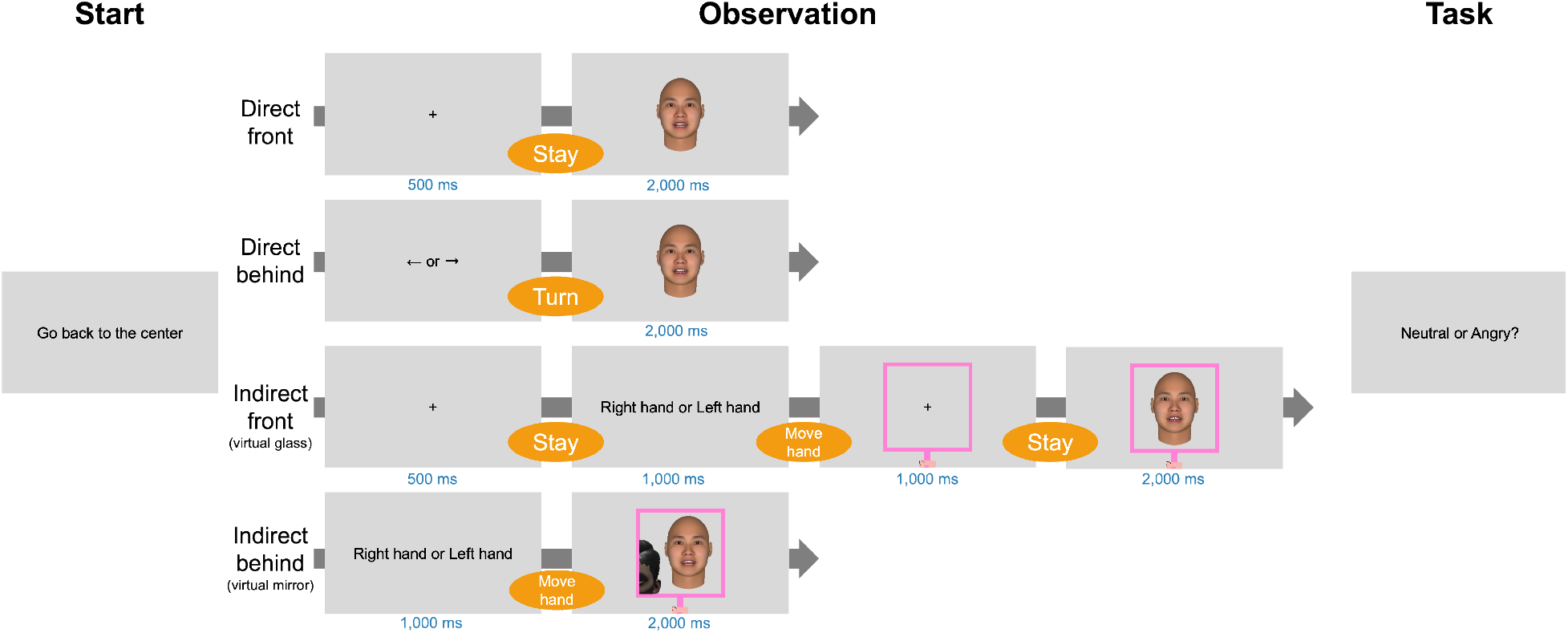
Procedure. The example stimulus shown here was not used in the experiments. Note that the background was not uniformly gray; instead, it consisted of a textured wall (see Procedure).

